# Disentangling the effects of hearing loss and age on amplitude modulation frequency selectivity

**DOI:** 10.1101/2023.07.15.549131

**Authors:** Jonathan Regev, Helia Relaño-Iborra, Johannes Zaar, Torsten Dau

## Abstract

The processing and perception of amplitude modulation (AM) in the auditory system reflect a frequency-selective process, often described as a modulation filterbank. Previous studies on perceptual AM masking reported similar results for older listeners with hearing impairment (HI) and young listeners with normal hearing (NH), suggesting no effects of age or hearing loss on AM frequency selectivity. However, recent evidence has shown that age, independently of hearing loss, adversely affects AM frequency selectivity. Hence, this study aimed to disentangle the effects of hearing loss and age. A simultaneous AM masking paradigm was employed, utilizing a sinusoidal carrier at 2.8 kHz, narrow-band noise modulation maskers, and target modulation frequencies of 4, 16, 64, and 128 Hz. The results obtained from young (n=3, 24-30 years) and older (n=10, 63-77 years) HI listeners were compared to previously obtained data from young and older NH listeners. Notably, the HI listeners generally exhibited lower (unmasked) AM detection thresholds and greater AM frequency selectivity than their NH counterparts in both age groups. Overall, the results suggest that age negatively affects AM frequency selectivity for both NH and HI listeners, while hearing loss improves AM detection and AM selectivity, likely due to the loss of peripheral compression.

## I. INTRODUCTION

Most sounds encountered in everyday environments are complex signals that exhibit dynamic fluctuations over time (Singh and Theunissen, 2003; Elliott and Theunissen, 2009). These fluctuations include variations in the temporal envelope, commonly referred to as amplitude modulation (AM). The auditory system demonstrates an acute sensitivity to AM, as evidenced by perceptual studies (e.g., Viemeister, 1979) and physiological investigations (see Joris *et al*., 2004 for a comprehensive review). Similar to spectral masking effects in the audio-frequency domain, masking effects also occur in the AM frequency domain, resulting in reduced sensitivity to a target AM in the presence of a masking AM (Bacon and Grantham, 1989; Houtgast, 1989; Dau *et al*., 1997a, 1997b; Ewert and Dau, 2000; Sek and Moore, 2003). Specifically, AM masking patterns provide evidence for a frequency-selective process, where the amount of AM masking decreases as the spectral distance between the masker and the target increases (Bacon and Grantham, 1989; Houtgast, 1989; Dau *et al*., 1997a; Ewert and Dau, 2000; Ewert *et al*., 2002; Sek and Moore, 2003; Moore *et al*., 2009; Sek *et al*., 2015; Füllgrabe *et al*., 2021a, 2021b; Conroy *et al*., 2023).

This frequency-selective characteristic of AM processing has been modelled using the concept of a modulation filterbank, based on the idea that AM fluctuations are decomposed through an array of relatively broad bandpass modulation filters with a constant quality (Q) factor of approximately 1-2 (e.g., Dau *et al*., 1997a, 1999; Ewert and Dau, 2000). Computational modelling studies have successfully applied the modulation filterbank concept to simulate data from various experimental paradigms, including simultaneous and non-simultaneous spectral and temporal signal detection and masking conditions (Dau *et al*., 1997a, 1997b, 1999; Verhey *et al*., 1999; Ewert and Dau, 2000; Ewert *et al*., 2002; Piechowiak *et al*., 2007; Jepsen *et al*., 2008; Jepsen and Dau, 2011; King *et al*., 2019), sound texture perception (McDermott and Simoncelli, 2011; McDermott *et al*., 2013; McWalter and Dau, 2015, 2017), auditory stream segregation (Elhilali *et al*., 2009; Christiansen *et al*., 2014), and speech intelligibility (Jørgensen and Dau, 2011; Jørgensen et al., 2013; Relaño-Iborra et al., 2016, 2019; Zaar and Dau, 2017; Zaar et al., 2017; Steinmetzger et al., 2019; Zaar and Carney, 2022; for a review, see Relaño-Iborra and Dau, 2022). Furthermore, the modulation filterbank is conceptually consistent with the temporal dimension of a ‘two-dimensional’ spectro-temporal modulation filterbank, inspired by neural responses to spectro-temporally varying stimuli in the auditory cortex of ferrets (Kowalski *et al*., 1996; Depireux *et al*., 2001) and supported by data from perceptual learning and masking conditions (Sabin *et al*., 2012; Oetjen and Verhey, 2015, 2017; Conroy *et al*., 2022), as well as models of speech intelligibility (Elhilali *et al*., 2003; Chi *et al*., 2005; Zilany and Bruce, 2007; Chabot-Leclerc *et al*., 2014). Overall, the versatility of the modulation filterbank model suggests that AM frequency selectivity is an essential auditory processing feature for quantitatively predicting perceptual data obtained with dynamically varying sounds. However, likely due to a lack of behavioral data, no modelling study was conducted investigating the effects of reducing the selectivity of the modulation filterbank, either due to age or hearing loss.

Several studies have investigated the effects of age and hearing loss on AM perception, yielding mixed results. In terms of age effects, some studies showed a significant age-related deterioration in AM detection (He *et al*., 2008; Füllgrabe *et al*., 2015; Wallaert *et al*., 2016), while other investigations did not find any effects on AM detection or AM depth discrimination (Schoof and Rosen, 2014; Paraouty *et al*., 2016; Schlittenlacher and Moore, 2016; Paraouty and Lorenzi, 2017). Studies comparing listeners with normal hearing to listeners with hearing impairment (referred to hereafter as NH and HI listeners, respectively) have found either similar or lower (i.e., better) AM detection thresholds for the HI listeners (e.g., Moore *et al*., 1992, 1996; Moore and Glasberg, 2001; Füllgrabe *et al*., 2003; Sek *et al*., 2015; Schlittenlacher and Moore, 2016; Wallaert *et al*., 2017; Wiinberg *et al*., 2019). The observed improvement in AM detection with hearing loss has been suggested to result from the loss of cochlear compression typically associated with sensorineural hearing loss (SNHL; Moore and Oxenham, 1998), which may lead to an increased internal representation of AM depth for HI listeners (Moore *et al*., 1996; Jennings *et al*., 2018). Additionally, the amplified neural coding of the envelope observed in mammals and humans with SNHL (Kale and Heinz, 2010, 2012; Henry *et al*., 2014; Zhong *et al*., 2014; Millman *et al*., 2017; Goossens *et al*., 2018; Decruy *et al*., 2020) may contribute, at least in part, to the effective increase in perceived AM depth with hearing loss. However, hearing loss has been shown to have a detrimental effect on supra-threshold AM depth discrimination (Schlittenlacher and Moore, 2016; Wiinberg *et al*., 2019), suggesting that the increase in perceived AM depth is not necessarily beneficial for AM depth discrimination.

Behavioral studies on AM frequency selectivity, where age and hearing loss were often confounded, generally did not yield conclusive evidence of an effect of hearing loss (Takahashi and Bacon, 1992; Lorenzi *et al*., 1997; Sek *et al*., 2015). These studies focused on masked-threshold patterns (MTPs), which assess the detection threshold of an AM target in relation to the (center) frequency of a masker of fixed modulation depth. Takahashi and Bacon (1992) compared MTPs (using noise carriers and sinusoidal modulation targets and maskers) from ten young NH listeners and three groups of ten older listeners with mild hearing loss (classified as ‘age-appropriate’ normal hearing). While a larger proportion of the older listeners could not complete some conditions of the AM masking task, no combined effect of age and hearing loss on AM frequency selectivity was found. Lorenzi *et al*. (1997) employed a similar approach with four young NH and three young HI listeners, finding comparable MTPs across the two groups, despite observations suggesting that HI listeners may experience a greater masking effect from modulation maskers located below the target modulation frequency. Additionally, Sek *et al*. (2015), using sinusoidal carriers and sinusoidal modulation maskers, found no differences in AM masking patterns between six young NH and nine older HI listeners. These results suggested that neither age nor hearing loss affect AM frequency selectivity (Sek *et al*., 2015).

However, recent results by Regev *et al*. (2023) showed a reduction in AM frequency selectivity associated with age when comparing young and older listeners with clinically normal hearing. These results are qualitatively consistent with the observations of reduced temporal selectivity inferred from auditory cortical functional magnetic resonance imaging (fMRI) responses by Erb *et al*. (2020). Regev *et al*. (2023), using sinusoidal carriers and noise modulation maskers, compared MTPs for eleven young NH listeners and ten older NH listeners (showing only a small mismatch in absolute hearing threshold at the test frequency of 10 dB on average; see Section IV in Regev *et al*., 2023 for an in-depth discussion). They found an age-related increase (i.e., deterioration) in AM detection thresholds and a decrease in AM frequency selectivity, with estimated modulation-filter Q factors that were approximately halved for the older NH listeners compared to the young NH listeners. The contrast between these recent findings and previous studies that did not identify a combined effect of age and hearing loss on AM frequency selectivity (Takahashi and Bacon, 1992; Sek *et al*., 2015) may have occurred because hearing loss and age have opposite effects with regard to AM frequency selectivity. In other words, hearing loss may ‘counteract’ the detrimental effects of ageing. However, a direct comparison of MTPs between young NH, older NH, and similarly aged older HI listeners, tested using the same experimental approach, has not yet been conducted.

The present study therefore aimed to independently evaluate the effects of hearing loss and age on AM frequency selectivity by collecting data from HI listeners who were approximately age-matched to the older NH listeners included in Regev *et al*. (2023), using the same experimental approach. Young HI listeners were also included to explore whether a perceptual ‘benefit’ from hearing loss could be observed in the absence of ageing effects. The study involved ten older and three young HI listeners, ranging in age from 63 to 77 and 24 to 30 years, respectively. MTPs were collected using an AM masking paradigm with sinusoidal carriers and bandpass Gaussian-noise modulation maskers (Ewert *et al*., 2002). Additionally, unmasked AM detection thresholds were obtained using sinusoidal carriers. The data obtained from the older HI listeners were compared to those obtained from the older NH listeners reported in Regev *et al*. (2023) to assess the effects of hearing loss on AM frequency selectivity. The data from the older HI listeners were also compared to those for the young NH listeners of Regev *et al*. (2023) to determine whether the lack of a combined effect of age and hearing loss observed in previous studies could be replicated (Takahashi and Bacon, 1992; Sek *et al*., 2015). The data reported in Regev *et al*. (2023) are publicly available from Regev *et al*. (2024). Despite the limited number of young HI listeners, their data were included to provide insight into the effects of hearing loss in the absence of ageing.

## II. METHODS

### A. Listeners

Ten older HI listeners, aged 63-77 years (mean age = 70.4 years, 2 female) and three young HI listeners, aged 24-30 years (mean age = 26 years, all female) participated in the study. On average, the older HI listeners were 4.5 years older than the older NH listeners tested in Regev *et al*. (2023), while the young HI listeners were less than one year older than the young NH listeners. All listening experiments were conducted monaurally. The individual audiograms for the test ears are shown in Figure 1. The older HI listeners had a high-frequency sloping hearing loss, classified as an N2 or N3 standard audiogram (Bisgaard *et al*., 2010).^1^ The young HI listeners had more heterogeneous audiograms. For each listener, the test ear was selected as the one where the audiometric threshold at 3 kHz was in the range 50 to 60 dB HL. These limits were chosen to ensure that the listeners had a significant hearing loss near the AM carrier frequency (2.8 kHz) and, at the same time, that the sound pressure level (SPL) would not exceed 85 dB, to avoid discomfort. All listeners had a SNHL, as indicated by air-bone gaps ≤ 15 dB.^2^ Working memory capacity was assessed with a reverse digit span test (RDS), implemented binaurally, as described in Fuglsang *et al*. (2020), with an added linear gain according to the Cambridge formula (CamEq; Moore and Glasberg, 1998) using a maximum-gain limit of 30 dB at any frequency. The young and older HI listeners were found to have similar average RDS scores (0.56 and 0.53 on a normalized scale, respectively), which are also comparable to the average RDS scores reported for the young NH listeners in Regev *et al*. (2023) and slightly, but not significantly, higher than for Regev *et al*.’s older NH cohort (0.59 and 0.42, respectively).

**Figure 1:**
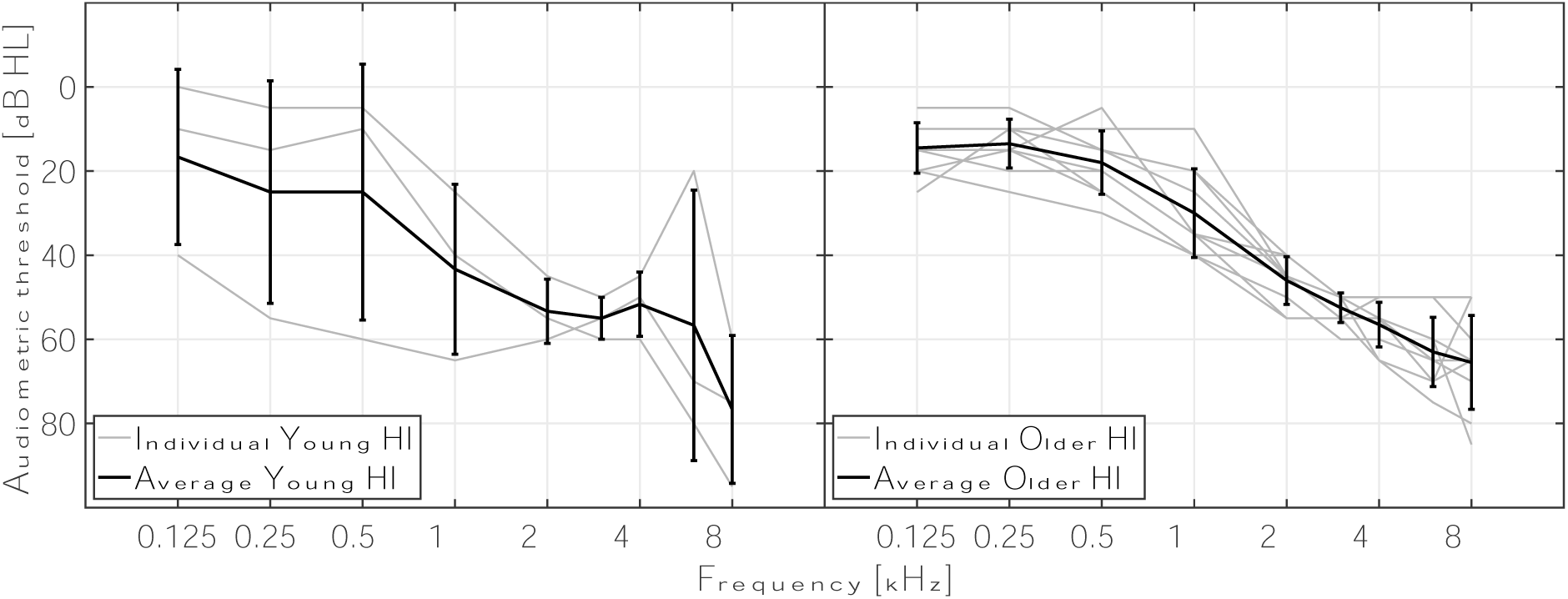
Individual and average audiograms for the test ear, for all participants. The left and right panels show the audiograms for the young and older HI listeners, respectively. Individual audiograms are shown in light gray, and group averages and standard deviations are shown in black.

All participants gave written informed consent. Ethical approval for the study was provided by the Science Ethics Committee of the Capital Region in Denmark (reference H-16036391). All participants were paid for their time.

### B. Procedure and apparatus

The experimental conditions and parameters were the same as described in Regev *et al*. (2023), with a few modifications to compensate for audibility loss. The stimulus presentation level in all experiments was adjusted to be 30 dB sensation level (SL) for all HI listeners, based on the hearing threshold at the carrier frequency, in an approach similar to that of Sek *et al*. (2015). This resulted in presentation levels of 78-80 dB SPL for the young HI listeners and 73-85 dB SPL for the older HI listeners (79 dB SPL on average for both groups).

All listening tests were conducted in double-walled soundproof booths. All thresholds were measured using a 3-interval 3-alternative forced-choice (3I-3AFC) paradigm with a 1-up 2-down tracking procedure, measuring the 70.7%-correct point on the psychometric function (Levitt, 1971). The experiments were implemented using the AFC toolbox in MATLAB (Ewert, 2013). The stimuli were digitally generated in MATLAB (The Mathworks Inc., 2015; Natick, Massachusetts) with a sampling frequency of 48 kHz, converted to analog using an RME Fireface soundcard (Haimhausen, Germany), and played through Sennheiser HDA200 headphones (Wedemark, Germany). The output level was calibrated using a GRAS RA0039 ear simulator (Holte, Denmark) and the frequency response of the headphones was compensated using an inverse digital filter to achieve a flat response at the tympanic membrane. The listeners received visual feedback during the tests (“correct”/“incorrect”). The order of testing was the same for both groups, as summarized in Appendix A.

The hearing threshold for the unmodulated pure-tone carrier at 2.8 kHz was measured adaptively. The signal had a duration of 300 ms, including 50-ms raised-cosine ramps, and the intervals were separated by 600 ms of silence. The step size was initially 8 dB, was then reduced to 4 dB after the first lower reversal, and was finally reduced to 2 dB after the second lower reversal. For any run, six reversals were obtained using the smallest step size and the threshold was defined as the average SPL of the target tone across them. The threshold measurement was repeated at least three times, and the final threshold was computed as the average across all repetitions.

### C. Experimental conditions

#### 1. Stimuli

The AM detection task and masked-threshold patterns (MTPs; Ewert and Dau, 2000) shared common aspects of the stimuli. For both tasks, the thresholds were measured for sinusoidal AM at target modulation frequencies of 4, 16, 64, and 128 Hz, imposed on a 2.8-kHz sinusoidal carrier. The carrier had a duration of 600 ms, including 50-ms raised-cosine onset and offset ramps. The target modulation (and masker modulation in the MTP task) had a duration of 500 ms, including 50-ms raised-cosine onset and offset ramps, and was temporally centered in the carrier. The target modulation was imposed with a 0-degree starting phase. The stimulus intervals were separated by 500 ms of silence. The stimuli for all three intervals were calibrated to the desired presentation level to ensure that no level cues were available. The tracking variable was the target’s modulation depth, expressed on a dB scale as 𝑀 = 20𝑙𝑜𝑔_10_(𝑚), where 𝑚 is the modulation depth on a linear scale. The step size was initially 4 dB, was reduced to 2 dB after the first lower reversal, and was finally reduced to 1 dB after the second lower reversal.

##### a. AM detection thresholds

For the AM detection task, the starting modulation depth of the target was −5 dB. For any run, six reversals were obtained using the smallest step size and the threshold was defined as the average target modulation depth across them. The measurement at each target modulation frequency was repeated three times, and the final threshold was defined as the average across repetitions. Additional measurements were included if the standard error across repetitions exceeded 2 dB, until that limit was met.^3^ All listeners completed two training runs (at the target modulation frequencies of 4 and 128 Hz). Figure 2 shows an example of the stimulus envelope for the reference interval (i.e., the unmodulated carrier, in panel A) and the target interval (i.e., the carrier modulated with the target modulation, in panel B) for a target modulation frequency of 128 Hz.

**Figure 2:**
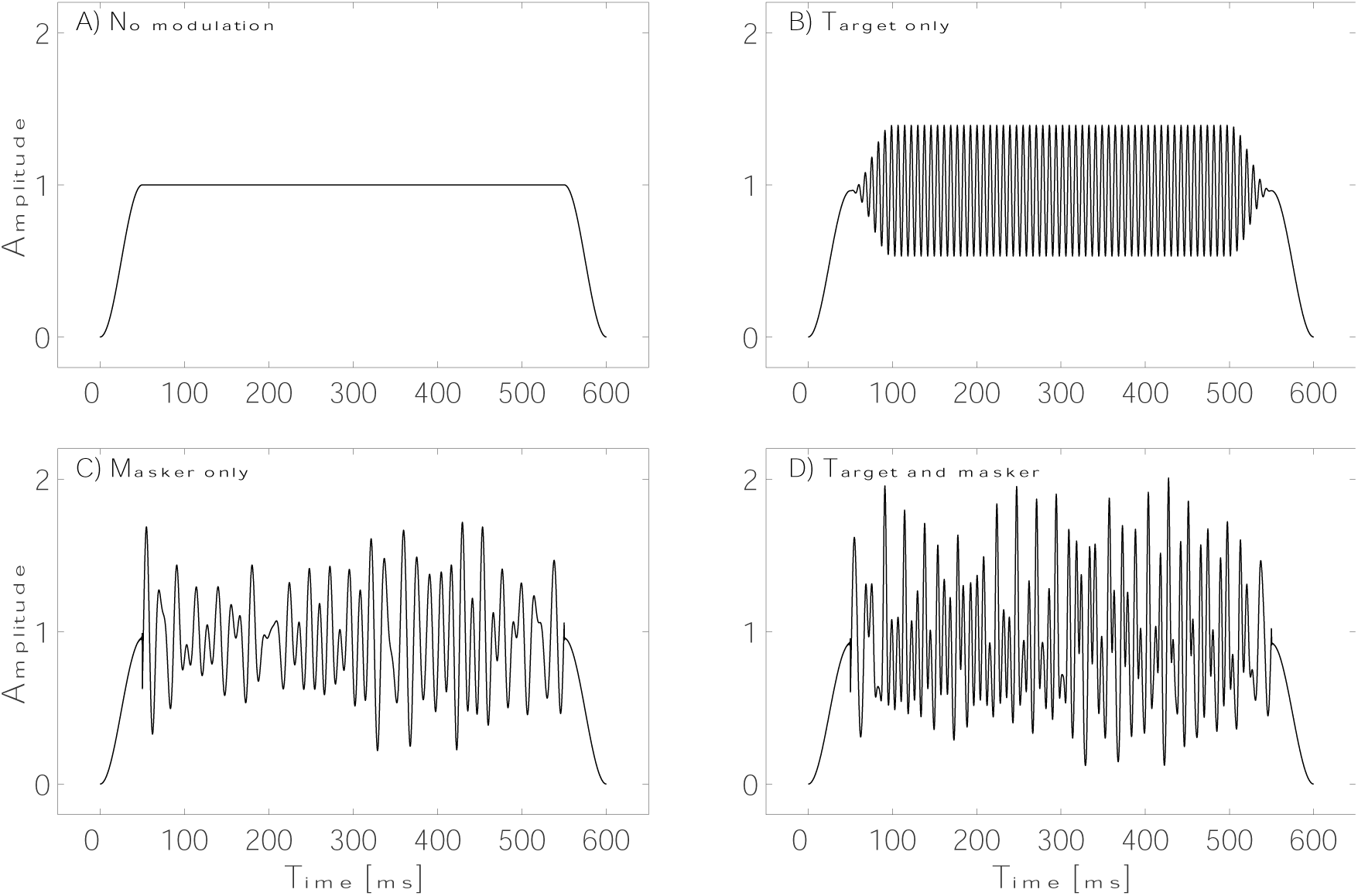
Examples of the temporal envelopes of the stimuli used in the AM detection and MTP experiments. Panel A: Reference interval in the AM detection task, represented by an unmodulated sinusoidal carrier (2.8 kHz). Panel B: Target interval in the AM detection experiment, containing the same carrier as in Panel A modulated by sinusoidal target modulation (128 Hz). Panel C: Reference interval in the MTP experiment, containing the same carrier as in Panel A modulated by narrowband-noise masker modulation (between 43.6 and 88.2 Hz, centered at 64 Hz). Panel D: Target interval in the MTP experiment, with the same carrier as in Panel A modulated by the sinusoidal target modulation and the narrowband-noise masker modulation.

##### b. Masked-threshold patterns

For the MTP task, the target and masker modulations were applied simultaneously to the carrier using a multiplicative approach, as described in Regev *et al*. (2023). The masker modulation was a narrow-band Gaussian noise with a fixed bandwidth corresponding to ½-octave when centered on the target modulation frequency. For target modulation frequencies of 4 and 16 Hz, the masker modulation was centered at frequencies ranging from −2 to +2 octaves relative to the target modulation frequency, in steps of 2/3 octaves. In order to avoid cues from spectrally resolved modulation sidebands, the highest masker-modulation center frequency was limited to 128 Hz, such that the highest masker-modulation center frequencies corresponded to +2/3 and 0 octaves for the target modulation frequencies of 64 and 128 Hz, respectively. For these target modulation frequencies, a masked threshold was also measured for masker modulations centered around 4 Hz, corresponding to −4 and −5 octaves relative to the target modulation frequencies, respectively. For these additional conditions, the lower cutoff frequency of the masker modulation was set to 1 Hz, thus effectively reducing the masker’s bandwidth. The resulting masker-modulation bandwidths and center frequencies for all conditions are summarized in Appendix B. For each measurement run, a 2-s long segment of the masker modulation was created. For each interval, a 500-ms segment, randomly cut out of the long masker modulation, was applied to the carrier. The root-mean square (rms) modulation depth of the masker modulation was fixed at −10 dB.

The starting modulation depth of the target was 0 dB. For any run, eight reversals were obtained using the smallest step size and the threshold was defined as the average target modulation depth across them. The measurement at each masker-modulation center frequency was repeated three times, and the final threshold was defined as the average across repetitions. To avoid overmodulation, the maximum allowed target modulation depth was 0 dB, and a safety check was introduced during the stimulus generation to minimize any potential residual overmodulation caused by the masker modulation. If the procedure required a target modulation depth above 0 dB more than three times over the course of a run, the run was stopped and an extra run was carried out to replace it. If this happened more than 5 times for any condition, the threshold was considered as unmeasurable. This was the case for a single older HI listener, whose thresholds could not be estimated for masker-modulation center frequencies of +2/3 and +4/3 octaves relative to the 4 Hz target modulation frequency. This listener’s masked thresholds at 4 Hz were therefore not included in the statistical analyses, or in the figures below, such that the MTP analyses at 4 Hz included only nine older HI listeners. Additional measurements were included if the standard error across repetitions exceeded 2 dB, until that limit was met.^4^ For each MTP, the listeners completed three training runs (at the masker-modulation center frequencies of −2, 0, and +2 octaves relative to the target modulation frequency). Figure 2 shows an example of the stimulus envelope for the reference interval (i.e., the carrier modulated with the masker modulation, in panel C) and the target interval (i.e., the carrier modulated with both the target and masker modulations, in panel D), for a target modulation frequency of 128 Hz and a masker-modulation center frequency of 64 Hz.

#### 2. Estimates of AM frequency selectivity

Ewert *et al*. (2002) and Regev *et al*. (2023) used the envelope power spectrum model of masking (EPSM; Ewert and Dau, 2000) to quantify the AM frequency selectivity reflected in their MTPs, by finding the Q-factor of the best-fitting bandpass modulation filter to the experimental data. However, high levels of asymmetry were observed in the MTPs in the present study (as described in detail below), which are not consistent with the EPSM’s assumption of symmetry (see section IV.D for a discussion of this aspect). Hence, a simplified approach was used here to quantify AM frequency selectivity. The best-fitting linear approximation (in terms of the least-squared error) was obtained separately for each skirt using the group-average masked thresholds for the masker locations ranging from −2 to 0 and 0 to +2 octaves. Each linear fit was constrained to cross the on-target masked threshold. The bandwidth of the asymmetrical modulation filter was then estimated by measuring the spectral distance between the 3-dB-down points on the fits obtained for each skirt when interpolating across masker locations ranging from −4 to +4 octaves. For the 128-Hz pattern, the high-frequency cutoff was estimated assuming symmetry around the target modulation frequency. Finally, the metric quantifying AM frequency selectivity was obtained as an overall ‘Q-factor’ by computing the ratio of the filter’s characteristic (modulation) frequency to the fitted bandwidth. The overall Q-factors describing the young NH and older NH average MTPs, obtained by Regev *et al*. (2023), were also re-evaluated using this linear approximation.

### D. Statistical analysis

Analyses of variances (ANOVAs) of linear mixed-effects model fittings were used to assess the differences between the results for the older HI listeners obtained in the present study and the data for the NH listeners obtained by Regev *et al*. (2023). Each model and corresponding ANOVA were applied twice. First, to estimate the effects of hearing loss in isolation from age effects, the results for the older HI listeners were compared with the results for the older NH listeners. Second, using an analysis comparable to that employed in previous studies, the results for the older HI listeners were compared with the results obtained for the young NH listeners. For the analysis of the AM detection thresholds, the listeners were considered as a random effect and group classification and modulation frequency were considered as fixed effects. The MTPs were analyzed separately for each target modulation frequency. The listeners were considered as a random effect and group classification and masker-modulation center frequency were considered as fixed effects. Levene’s test was used to assess the homoskedasticity of the residuals and Shapiro-Wilk’s test was used to assess the normality of the residual’s distribution.^5^ Post-hoc analyses were conducted using estimated marginal means and a Holm-Bonferroni correction criterion was applied for multiple comparisons. The level of statistical significance was set to 5% in all analyses. Due to the low number of young HI listeners (3), no statistical analysis was performed to compare their data to those of the other groups. This limitation is further discussed in section IV.D.

## III. RESULTS

### A. AM detection thresholds

Figure 3 shows the mean AM detection thresholds as a function of modulation frequency for the young HI (diamonds), older HI listeners (circles), young NH (squares), and older NH listeners (triangles), the latter two datasets from Regev *et al*. (2023). Substantial levels of individual variability were observed (see supplementary Figure 1). The statistical analysis comparing the thresholds for the older HI and the older NH listeners revealed a significant main effect of group [𝐹_1,18_ = 21.21, 𝑝 < 0.001] and modulation frequency [𝐹_3,54_ = 22.84, 𝑝 < 0.001]. The post-hoc analysis revealed that the older HI listeners had significantly lower (i.e., better) AM detection thresholds than the older NH listeners at all modulation frequencies [effect size (expressed as difference in 𝑀; Δ𝑀) = −6.0 dB, standard error (𝑆𝐸) = 2.0 dB; Δ𝑀 = −8.5 dB, 𝑆𝐸 = 2.0 dB; *Δ*𝑀 = −7.4 dB, 𝑆𝐸 = 2.0 dB; *Δ*𝑀 = −7.0 dB, 𝑆𝐸 = 2.0 dB, for 4, 16, 64, and 128 Hz, respectively].

**Figure 3.**
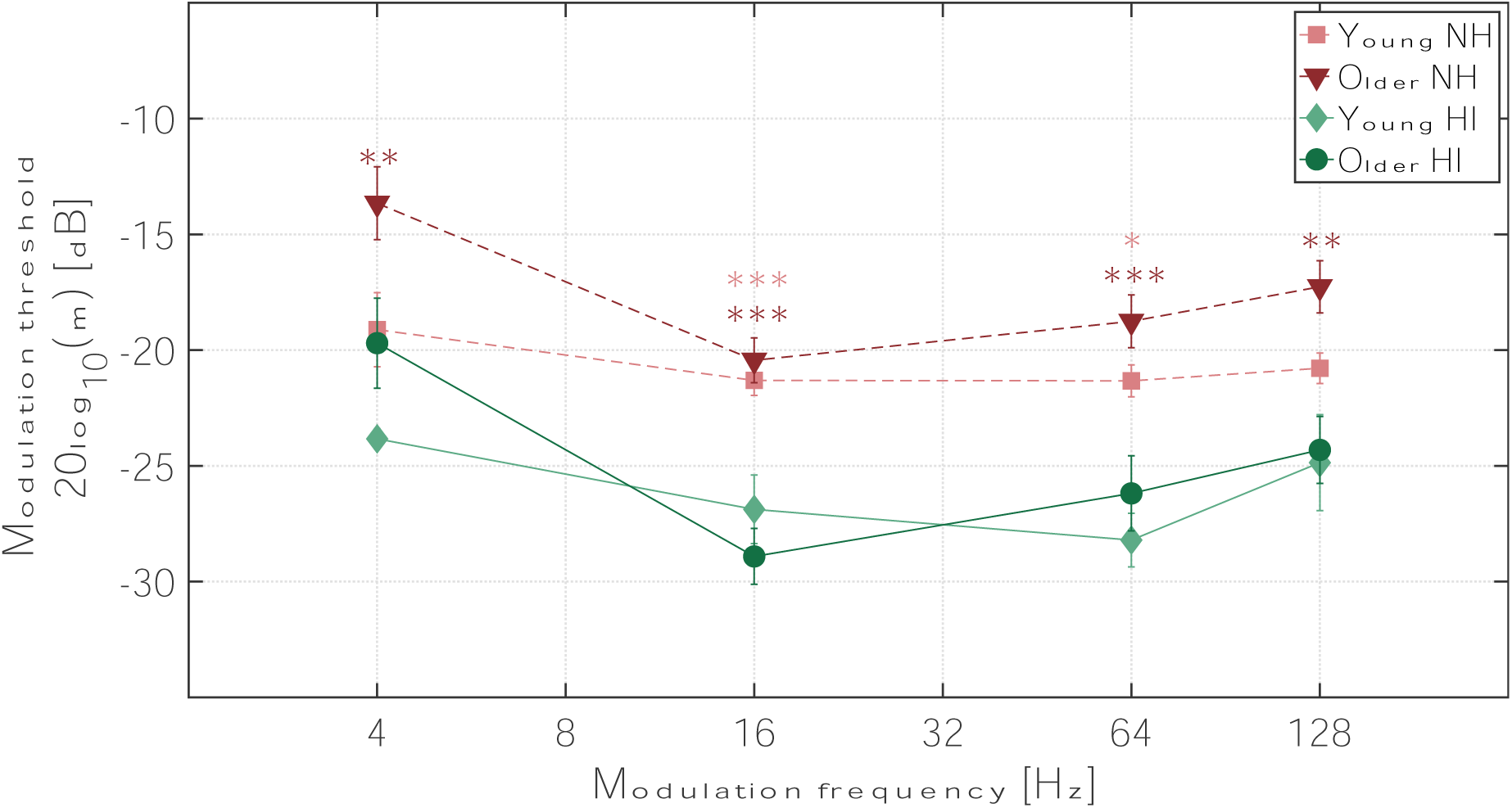
(Color online): Average AM detection thresholds for each listener group as a function of modulation frequency. The data for the young HI listeners are indicated by diamonds connected by solid lines. The data for the older HI listeners are shown as circles connected by solid lines. The results for the young and the older NH listeners (dashed lines; data from Regev et al., 2023) are shown as triangles and squares, respectively. Means and standard errors are shown. The stars indicate the level of statistical significance of the differences in thresholds between the older HI and older NH listeners (dark stars), and the older HI and young NH listeners (light stars; *\* p < 0.05, ** p < 0.01, *** p < 0.001*).

The statistical analysis comparing the thresholds for the older HI and the young NH listeners revealed a significant main effect of group [𝐹_1,19_ = 8.13, 𝑝 = 0.01] and modulation frequency [𝐹_3,57_ = 14.66, 𝑝 < 0.001], and a significant interaction [𝐹_3,57_ = 5.26, 𝑝 = 0.003]. The post-hoc analysis revealed that the older HI listeners had significantly lower AM detection thresholds than the young NH listeners for modulation frequencies of 16 Hz [Δ𝑀 = −7.6 dB, 𝑆𝐸 = 1.8 dB] and 64 Hz [*Δ*𝑀 = −4.9 dB, 𝑆𝐸 = 1.8 dB].

### B. Masked-threshold patterns (MTPs)

Figure 4 shows the mean MTPs for the older HI (circles) and the older NH listeners (triangles; data from Regev *et al*., 2023). Each panel shows the MTPs for a specific target modulation frequency. The mean AM detection thresholds are indicated by the open symbols (replotted from Figure 3). The statistical analysis comparing the older HI and the older NH listeners revealed a significant main effect of group for all target modulation frequencies [𝐹_1,17_ = 7.31, 𝑝 = 0.015; 𝐹_1,18_ = 20.18, 𝑝 < 0.001; 𝐹_1,18_ = 8.70, 𝑝 = 0.009; 𝐹_1,18_ = 14.43, 𝑝 = 0.001, for 4, 16, 64, and 128 Hz, respectively]. A significant interaction between listener group and masker-modulation center frequency was found for the target modulation frequencies of 4 Hz [𝐹_6,102_ = 3.36, 𝑝 = 0.005] and 16 Hz [𝐹_6,108_ = 4.64, 𝑝 < 0.001], whereas the interaction approached, but did not reach, significance for the target modulation frequency of 64 Hz [𝐹_5,90_ = 2.01, 𝑝 = 0.085]. For the target modulation frequency of 4 Hz, the post-hoc analysis revealed that the older HI listeners had significantly lower (better) masked thresholds than the older NH listeners for the masker modulations centered at −2, −4/3, and −2/3 octaves relative to the target frequency. At 16 Hz, the post-hoc analysis revealed that the older HI listeners had significantly lower masked thresholds than the older NH listeners for the masker modulations centered at −2, −4/3, and +2 octaves relative to the target frequency. At 64 Hz, the post-hoc analysis revealed that the older HI listeners had significantly lower masked thresholds than the older NH listeners for the masker modulations centered at −4, −2, and +2/3 octaves relative to the target frequency. Finally, at 128 Hz, the post-hoc analysis revealed that the older HI listeners had significantly lower masked thresholds than the older NH listeners for the masker modulations centered at −5, −2, − 4/3, and 0 octaves relative to the target frequency. The details of the post-hoc analyses can be found in Appendix B. Overall, the MTPs were sharper for the older HI listeners than for the older NH listeners, for the target modulation frequencies of 4 Hz (on the low-frequency skirt), 16 Hz, and 64 Hz. Conversely, the high-frequency skirts of the 4-Hz MTPs suggests similar selectivity for the older HI and NH listeners. Finally, the MTP at 128 Hz reflects a general improvement (i.e., decrease) of the masked thresholds for the older HI listeners, but an overall broader pattern, compared to that for the older NH listeners, as shown by the fitted Q-factors displayed in Table 1 and discussed further below. High levels of asymmetry were also observed for the older HI listener’s MTPs, as quantified by the slopes of the obtained linear fits (see supplementary Figure 2). At the target modulation frequency of 4 Hz, this was reflected by a steeper slope of the low-frequency skirt (3 dB/octave) than the high-frequency skirt (1 dB/octave) and an off-target peak threshold (for the masker centered at +2/3 octaves relative to the target frequency). A similar effect was found at 16 Hz, with a steeper slope of the low-frequency skirt (5.6 dB/octave) than of the high-frequency skirt (4.6 dB/octave). At 64 Hz, the older HI listeners gave a much shallower slope of the low-frequency skirt (3.6 dB/octave) than of the high-frequency skirt (9.3 dB/octave), although the latter was computed based only on the average masked threshold at +2/3 octaves.

**Figure 4.**
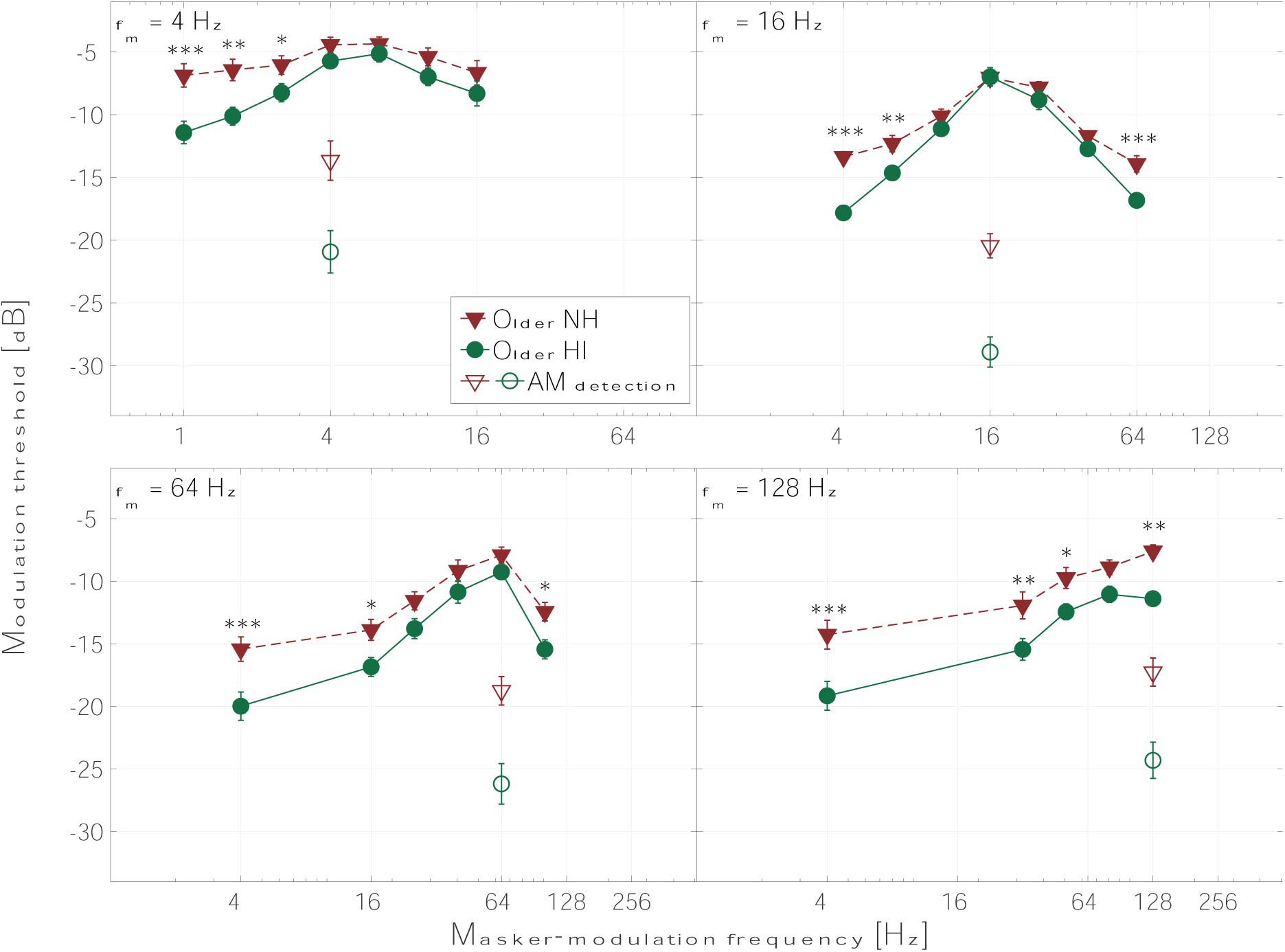
(Color online): Average MTPs (filled symbols) and unmasked AM detection thresholds (open symbols, replotted from Figure 3) for the older HI and the older NH listeners. The data for the HI listeners are indicated by the circles connected by solid lines and the data for the NH listeners (from Regev *et al*., 2023) are shown as triangles connected by dashed lines. Each panel represents the results for one target modulation frequency: 4 Hz (top left), 16 Hz (top right), 64 Hz (bottom left), and 128 Hz (bottom right). The data from one of the older HI listeners were excluded from the 4-Hz panel for both the MTP and AM detection threshold. Means and standard errors are shown. Stars indicate the level of statistical significance of the differences in masked thresholds between the two groups (*\* p < 0.05, ** p < 0.01, *** p < 0.001*).

**Table 1:** Overall fitted Q-factors, defined as the ratio of the filter’s characteristic frequency to its bandwidth estimated at the 3-dB-down point, and MSEs for the modulation filters fitted to the average group data using an asymmetric linear fitting approach. The table also shows the ratio of Q-factors for each group to the Q-factor for the young NH listeners

| Target<br>modulation<br>frequency [Hz] | Estimated Q-factor<br>(Overall MSE [dB]) |  |  |  | Q/Q<sub>YNH</sub> |  |  |
| --- | --- | --- | --- | --- | --- | --- | --- |
|  | YNH | ONH | YHI | OHI | ONH | YHI | OHI |
| 4 | 0.77<br>(0.35) | 0.10<br>(0.16) | 0.81<br>(0.78) | 0.14<br>(0.30) | 0.13 | 1.05 | 0.18 |
| 16 | 0.96<br>(0.13) | 0.75<br>(0.44) | 1.24<br>(0.38) | 1.13<br>(0.30) | 0.78 | 1.29 | 1.18 |
| 64 | 1.38<br>(0.15) | 1.14<br>(0.09) | 1.90<br>(1.0) | 1.45<br>(0.14) | 0.83 | 1.38 | 1.05 |
| 128 | 0.96<br>(0.05) | 0.40<br>(0.10) | 0.82<br>(0.98) | 0.27<br>(0.96) | 0.42 | 0.85 | 0.28 |

Figure 5 shows the mean MTPs for the young HI (diamonds) and older HI listeners (circles) and the data from the young NH listeners (squares) obtained by Regev *et al*. (2023). Each panel shows the MTPs for a specific target modulation frequency. The mean AM detection thresholds are also shown and indicated by the open symbols (replotted from Figure 3). The statistical analysis comparing the older HI and the young NH listeners showed no main effect of group for any target modulation frequency. However, there was a significant interaction between listener group and masker-modulation center frequency for all target modulation frequencies [𝐹_6,108_ = 3.95, 𝑝 = 0.001; 𝐹_6,114_ = 8.1, 𝑝 < 0.001; 𝐹_5,95_ = 4.12, 𝑝 = 0.002; 𝐹_4,76_ = 6.3, 𝑝 < 0.001, for 4, 16, 64, and 128

**Figure 5.**
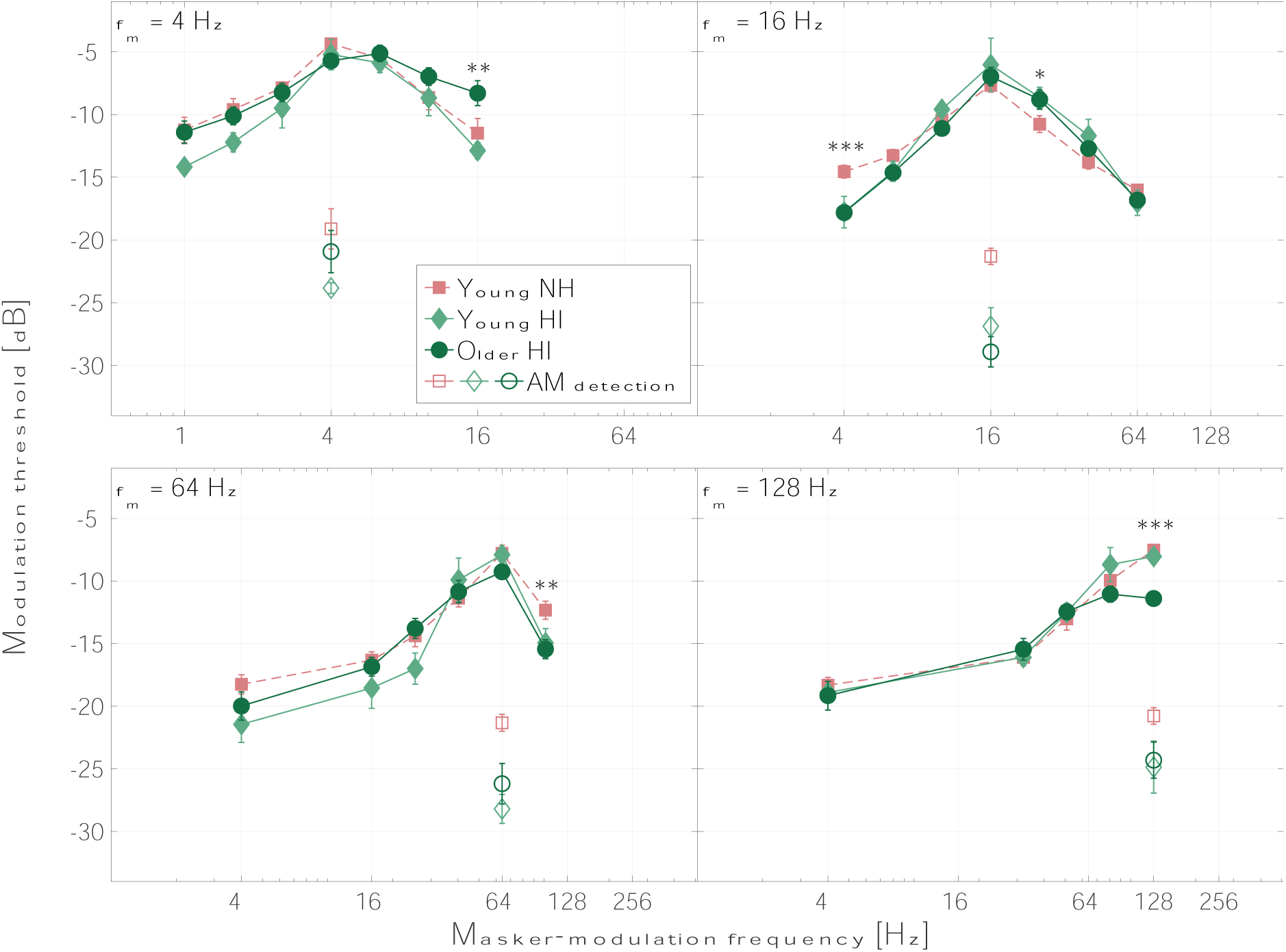
(Color online): Average MTPs (filled symbols) and unmasked AM detection thresholds (open symbols, replotted from Figure 3). The data for the young and older HI listeners (solid lines) are indicated by the diamonds and circles, respectively. The data for the young NH listeners are shown as dashed lines and squares (data from Regev *et al*., 2023). Each panel represents the results for one target modulation frequency: 4 Hz (top left), 16 Hz (top right), 64 Hz (bottom left), and 128 Hz (bottom right). The data from one of the older HI listeners were excluded from the 4-Hz panel for both the MTP and AM detection thresholds. Means and standard errors are shown. Stars indicate the level of statistical significance of the differences in masked thresholds between the young NH and older HI listeners (*\* p < 0.05, ** p < 0.01, *** p < 0.001*).

Hz, respectively]. For the target modulation frequency of 4 Hz, the post-hoc analysis showed that the older HI listeners had significantly higher (worse) masked thresholds than the young NH listeners for the masker centered at +2 octaves relative to the target. At 16 Hz, the older HI listeners had significantly lower (i.e., better) masked thresholds for the masker centered at −2 octaves relative to the target frequency and significantly higher masked thresholds than the young NH listeners for the masker centered at +2/3 octaves above the target frequency. At 64 Hz, the older HI listeners had significantly lower masked thresholds than the young NH listeners for the masker centered at +2/3 octaves above the target frequency. Finally, at 128 Hz, the older HI listeners had significantly lower masked thresholds than the young NH listeners for the masker centered at the target modulation frequency. Details of the post-hoc analyses can be found in Appendix B.

Although no statistical analysis including the data for the young HI listeners was performed, due to the low number of participants, their results provide insights into the effects of hearing loss in the absence of ageing. Similar to the MTPs for the older HI listeners, high levels of asymmetry were observed in the MTPs of the young HI listeners. At the target modulation frequency of 4 Hz, this was reflected by a steeper slope of the low-frequency skirt (4.8 dB/octave) than the high-frequency skirt (3.3 dB/octave). At 16 Hz, the low-frequency skirt (6 dB/octave) was again steeper than the high-frequency skirt (5 dB/octave). At 64 Hz, the young HI listeners displayed a much shallower low-frequency skirt (5.6 dB/octave) than high-frequency skirt (10.6 dB/octave), although the latter was computed based only on the average masked threshold at +2/3 octaves.

Table 1 summarizes the overall Q-factors fitted to the average data for each listener group and shows the mean square errors (MSEs, in dB) of the fit in parentheses. The table also shows the ratio of the Q-factors for each group to the Q-factor for the young NH listeners. A ratio higher than 1 indicates sharper tuning for the listener group in question relative to the young NH listeners. For the older NH listeners, the reduced Q-factor ratio reflects the age-related reduction of AM frequency selectivity shown by Regev *et al*. (2023). In contrast, the results for the older HI listeners suggest sharper tuning for this group than for the older NH listeners for the modulation frequencies of 4, 16, and 64 Hz, and reduced AM frequency selectivity at 128 Hz. Compared to the young NH listeners, the older HI listeners show similar, or slightly greater, AM frequency selectivity for the modulation frequencies of 16 and 64 Hz, and broader tuning at 4 and 128 Hz. The young HI listeners’ results suggest generally greater AM frequency selectivity than for the older HI listeners, and even higher Q-factors (i.e., sharper tuning) than the young NH listeners for the modulation frequencies of 4, 16, and 64 Hz.

## IV. DISCUSSION

### A. Summary of main findings

This study aimed to investigate the potentially opposite effects of hearing loss and age on AM frequency selectivity. These opposing effects were suggested based on recent findings of an age-related reduction in AM frequency selectivity (Regev et al., 2023), which appeared to contradict previous conclusions that neither age nor hearing loss affect AM frequency selectivity (Takahashi and Bacon, 1992; Sek *et al*., 2015). Behavioral MTPs and unmasked AM detection thresholds were collected from three young and ten older HI listeners (24-30 and 63-77 years). The results were compared to the data obtained from young and older NH listeners by Regev *et al*. (2023), publicly available from Regev *et al*. (2024). Both HI groups showed lower AM detection thresholds than their similarly aged NH counterparts, and the older HI listeners also showed lower detection thresholds than the young NH listeners for target modulation frequencies above 4 Hz (Figure 3). Furthermore, for target modulation frequencies below 128 Hz, the older HI listeners generally showed greater AM frequency selectivity than the older NH listeners (Figure 4). When compared to the young NH listeners, the older HI listeners displayed an asymmetric broadening of the MTP at the target modulation frequency of 4 Hz (with a shallower high-frequency skirt), but similar or even slightly sharper tuning at 16 and 64 Hz (Figure 5). At the target modulation frequency of 128 Hz, the older HI listeners showed lower masked thresholds overall than the older NH listeners (with an average offset of 3.4 dB) and similar masked thresholds to the young NH listeners for the off-target maskers, effectively indicating reduced AM frequency selectivity compared to both NH groups. The data from the young HI listeners suggested greater AM frequency selectivity than the older HI listeners, and generally sharper tuning than the young NH listeners (as can be seen in Figure 5). Notably, both the young and older HI listeners showed high levels of asymmetry in their MTPs, with generally steeper low-frequency than high-frequency skirts. Finally, the best-fitting Q-factor supported previous observations suggesting that AM frequency selectivity for HI listeners is similar to, or greater than, that for similarly aged NH listeners, except for the highest target modulation rate of 128 Hz (Table 1).

### B. Opposite effects of hearing loss and age

The results for both the AM detection and MTP tasks suggest a general perceptual benefit from hearing loss, in terms of increased detectability of the target modulation and a sharpening of the masked patterns. This benefit effectively counteracts the detrimental effects of age on AM detection and AM frequency selectivity. The improvement of AM detection thresholds for HI listeners compared to NH listeners is consistent with previous studies (e.g., Bacon and Gleitman, 1992; Moore and Glasberg, 2001; Sek *et al*., 2015; Schlittenlacher and Moore, 2016; Wallaert *et al*., 2017; Wiinberg *et al*., 2019). It has been suggested that this improvement is due to an effectively increased internal representation of the signal’s AM depth resulting from the loss of compression commonly associated with SNHL (Moore *et al*., 1996; Jennings *et al*., 2018). The trends observed for the young HI listeners support the notion that greater age is detrimental to AM detection whereas SNHL provides a benefit. Previous AM masking studies comparing young NH and older HI listeners using low target modulation frequencies (≤ 16 Hz) found no differences in MTPs (Takahashi and Bacon, 1992; Sek *et al*., 2015) and concluded that neither SNHL nor age affect AM frequency selectivity. The present results are broadly consistent with the observation that MTPs are similar for young NH and older HI listeners for low modulation frequencies, although an asymmetric broadening of the 4-Hz MTP was found for the older HI listeners, with a shallower high-frequency skirt. However, the data also showed that SNHL generally improves AM frequency selectivity for listeners of similar age for target modulation frequencies below 128 Hz. In contrast, at 128 Hz, the masked thresholds were lower for the older HI listeners than for the older NH listeners and effectively indicated a reduction in AM frequency selectivity for the older HI listeners. Additionally, a slight broadening of the pattern at 128 Hz was observed for young HI listeners compared to the young NH listeners.

The results of the present study show that young HI listeners generally perform better than older HI listeners, as shown by Regev *et al*. (2023) for NH listeners. Furthermore, although a sharpening of the MTPs for HI listeners was not observed on both skirts of the patterns, HI listeners generally outperformed similarly aged NH listeners in terms of AM detection thresholds and tuning of the MTPs. Factors that may have influenced these results, such as the difference in presentation levels across the NH and HI listeners, are discussed in section IV.D.

### C. Potential mechanism underlying the perceptual benefit of hearing loss

The underlying auditory processes that account for the sharper MTPs for the older HI listeners than for the older NH listeners may not be straightforward. It seems unlikely that SNHL would improve AM frequency selectivity by ‘restoring’ the bandwidth of the hypothetical modulation filters. Instead, the greater AM frequency selectivity observed here for the HI listeners may result from the reduced peripheral compression that is typically associated with SNHL. One notable characteristic of the data for the HI listeners is the strong asymmetry of the MTPs, with a steeper slope on the low-frequency than the high-frequency skirt. This was also found by Regev *et al*. (2023) for their NH listeners, although the effect was less prominent than in the present study. The asymmetry suggests that an additional cue for the detection of the target may be available when the modulation maskers are centered below - but not above - the target modulation frequency. The existence of such a cue was discussed in previous studies by Bacon and Grantham (1989) and Strickland and Viemeister (1996), who found negative masking when an AM masker was centered below the target modulation frequency. They argued that listeners may be able to detect the target modulation in the temporal dips of the masker modulation, or ‘listen in the dips’, which is a detection cue analogous to the release from masking provided by envelope fluctuations in spectral masking tasks (e.g., Buus, 1985). In the present study, the random dips occurring in the modulation masker would be longer than (or close to) one cycle of the target for maskers centered below the target, while this would not be the case for modulation maskers centered above the target. It is possible that these local temporal features aid the detection of the target in the presence of low-frequency maskers. The data suggest that such a cue, if present, may be more salient for HI listeners than for NH listeners. The loss of cochlear compression for HI listeners might increase the target’s AM depth glimpsed in the temporal valleys of the masker, further enhancing the detectability of the target. This notion is consistent with the findings of Wallaert *et al*. (2017), which showed a greater improvement in AM detection thresholds for HI listeners than for NH listeners when increasing the numbers of cycles of the target modulation, suggesting enhanced temporal integration for the HI listeners. This may imply that HI listeners require fewer cycles of the target modulation to detect its presence, potentially contributing to the improved performance measured on the lower skirt of the MTPs.

If this detection mechanism indeed mediated the sharper MTPs for HI listeners, investigating AM frequency selectivity using an AM forward-masking paradigm should eliminate the perceptual benefit of SNHL. AM forward masking has been used as an alternative approach for investigating AM frequency selectivity (Wojtczak and Viemeister, 2005; Moore *et al*., 2009; Füllgrabe *et al*., 2021a, 2021b) because it avoids extraneous cues resulting from the interaction of the target and masker modulation. While it remains unclear whether AM simultaneous and AM forward-masking paradigms reveal the same underlying AM frequency-selective process, a study comparing age-matched NH and HI listeners using such a paradigm could provide valuable insights into the origins of the observed asymmetry. However, such a study has not been conducted thus far. Additionally, computational auditory models could be employed to assess the plausibility of ‘listening in the dips’ as a mechanism underlying the present results.

The hypothesis that increased sensitivity to the temporal envelope, resulting from the loss of cochlear compression, offers a perceptual advantage for HI listeners is supported by the findings of Bianchi *et al*. (2016). They demonstrated that fundamental-frequency discrimination of unresolved complex tones was comparable for older HI and young NH listeners when the reduced cochlear compression resulted in an effective amplification of the modulation power of the stimulus (i.e., when the envelope was maximally peaky), while performance was worse for the older HI listeners when the envelope was much flatter and hence largely unaffected. Partially connected with the loss of cochlear compression, the better-than-normal AM detection thresholds and AM frequency selectivity observed here for the HI listeners may be related to the increased neural coding of the envelope which has been shown both in animal models and humans (Kale and Heinz, 2010, 2012; Henry *et al*., 2014; Zhong *et al*., 2014; Millman *et al*., 2017; Goossens *et al*., 2018; Decruy *et al*., 2020). This increased internal representation of the envelope, while it may reflect a listening advantage for simple AM detection and masking paradigms such as the ones considered in the present study, has been suggested ultimately to be detrimental to the intelligibility of speech in noise (Millman *et al*., 2017; Goossens *et al*., 2018; Decruy *et al*., 2020). Hence, it could potentially contribute to the difficulties experienced by HI listeners in complex listening scenarios. Similar conclusions were drawn by studies reporting a connection between poorer speech intelligibility and increased neural coding of the envelope in older NH listeners (e.g., Goossens *et al*., 2018; Decruy *et al*., 2019).

### D. Limitations of the study

The presentation levels used in the present study were not directly comparable to those employed by Regev *et al*. (2023) with NH listeners. Regev *et al*. used a fixed level of 65 dB SPL (corresponding to average sensation levels of 67 and 57 dB SL for the young and older NH listeners, respectively), while the present study adjusted the level for the HI listeners to be 30 dB SL (corresponding to 79 dB SPL on average), following a similar approach to Sek *et al*. (2015). This methodological difference implies that the presentation level was both lower in terms of SL (by 27-37 dB) and higher in terms of SPL (by 14 dB) for the HI listeners than for the NH listeners. Previous studies have highlighted the effect of the interaction between presentation level and hearing status on AM detection thresholds, often finding lower AM detection thresholds for HI listeners than for NH listeners at equal SL, but similar thresholds at equal SPL (Bacon and Gleitman, 1992; Moore and Glasberg, 2001; Füllgrabe *et al*., 2003; Schlittenlacher and Moore, 2016; Wallaert *et al*., 2017; Moore *et al*., 2019), although some studies still found differences at equal SPL (Sek *et al*., 2015; Jennings *et al*., 2018). Hence, it is possible that the present results were influenced by the difference in presentation levels between the NH and HI listeners. However, the present finding that the HI listeners showed lower AM detection thresholds than the NH listeners at both lower SLs and higher SPLs seems consistent with previous studies that reported similar results (Füllgrabe *et al*., 2003; Jennings *et al*., 2018). Füllgrabe *et al*. (2003) observed an increased difference in AM detection thresholds between NH and HI listeners when the experiment was repeated with a fixed SL for all participants, relative to the condition where the HI listeners were tested with a higher SPL and lower SL than the NH listeners. Additionally, Wojtczak (2011) investigated the effects of carrier level on AM frequency selectivity for NH listeners and found that a reduction in SL led to an increase in AM detection thresholds and a broadening of the MTPs. Therefore, it is likely that testing NH and HI listeners at equal SL would have further increased the observed differences between the two groups in the present study. However, it remains unclear whether lower AM detection thresholds and sharper MTPs for the HI listeners than for the NH listeners would still be found if they were tested at equal SPL.

It is plausible that the underlying pathology of the hearing loss was different for the young and older HI listeners, potentially affecting the effects on sound perception. Although all HI listeners had a SNHL, the origin of the hearing loss in the young HI listeners was unfortunately unknown. Additionally, the small number of young HI listeners included in this study, along with the significant variability in their audiograms (except for the frequency of interest), limited the conclusions that could be drawn from their data. In spite of these aspects, the results for the young HI listeners in this study may offer some valuable insights. Indeed, the data obtained across the three young HI listeners were consistent with the observations made based on the other listener groups. Specifically, young listeners exhibited sharper tuning than older listeners with the same hearing status, and HI listeners generally demonstrated sharper tuning than similarly aged NH listeners. Conducting further investigations with a larger sample of young HI listeners may help validate the trends identified in the present study.

Finally, the method used to quantify AM frequency selectivity in the present study was limited by the pronounced asymmetry of the MTPs. Previous studies quantified the tuning of the hypothetical modulation filters using the EPSM (Ewert and Dau, 2000; Ewert *et al*., 2002; Regev *et al*., 2023), which assumes symmetrical filters centered around the characteristic modulation frequency. However, the present results do not support this assumption, indicating a need to revise the EPSM to incorporate more flexible filter shapes. Similar conclusions were drawn by Regev *et al*. (2023), who observed large variability in individual MTPs. Consequently, the Q-factor values reported in this study for the young and older NH listeners differ from those reported by Regev *et al*. (2023) due to the different approaches used to quantify AM frequency selectivity. However, the overall finding of lower Q factors for the older NH listeners than for the young NH listeners is consistent between the two investigations.

## V. CONCLUSIONS

This study investigated the effects of hearing loss on AM frequency selectivity by employing a simultaneous AM masking task. Ten older and three young HI listeners were tested, ranging in age from 63-77 and 24-30 years, respectively. These age ranges were approximately matched to those of the young and older NH listeners tested by Regev *et al*. (2023). Although no statistical analysis including the young HI data could be conducted, due to the limited number of young HI listeners, their results provide valuable insight into the effects of hearing loss in the absence of ageing. The HI listeners showed lower AM detection thresholds and generally sharper MTPs than the NH listeners, suggesting that hearing loss may counteract the detrimental effects of age on AM frequency selectivity. The MTPs showed strong asymmetry, with generally steeper low-frequency skirts. A potential mechanism for explaining these results is the detection of the target modulation in the dips of low-frequency masker modulations, coupled with the effective increase in AM depth resulting from the loss of cochlear compression. This suggests that the observed sharpening of the MTPs for HI listeners may be linked to the loss of cochlear compression rather than a ‘restoration’ of the bandwidth of the underlying hypothetical modulation filters. Differences in presentation level between the HI and NH listeners may have influenced the threshold differences, so the results of this study motivate further investigations into the effects of hearing loss on AM masking and AM frequency selectivity, as well as their interaction with age-related effects. In addition to the loss of cochlear compression, the lower AM detection thresholds and greater AM frequency selectivity observed in HI listeners may be associated with increased neural coding of the envelope, which has been suggested to be detrimental to speech intelligibility in complex listening scenarios. Hence, future studies examining the effects of hearing loss on AM masking can offer valuable insights into the underlying mechanisms that contribute to the perceptual benefits observed for HI listeners and their implications for complex auditory perception and speech intelligibility.

## ACKNOWLEDGMENTS

The authors thank Borgný Súsonnudóttir Hansen for her contribution to the data collection, as well as Erik Kjærbøl and Jesper Borchorst Yde for their precious help in recruiting young HI participants. We also thank Stephan Ewert for helpful suggestions and for kindly sharing experimental details and data from earlier publications, Jens Hjortkjær and Jonatan Märcher-Rørsted for kindly providing their implementations of the reverse digit span test and of the Cambridge formula (CamEQ; Moore and Glasberg, 1998), as well as Raul Sanchez-Lopez and Beverly Wright for helpful insights and discussions. We are thankful to Brian C. J. Moore and two other anonymous reviewers for their insightful comments during the review process.This study was supported by the Oticon Centre of Excellence for Hearing and Speech Sciences (CHeSS).

## AUTHOR DECLARATIONS

### Ethics Approval

All participants gave written informed consent and were financially compensated for their time. Ethical approval for the study was provided by the Science Ethics Committee of the Capital Region in Denmark (reference H-16036391).

## DATA AVAILABILITY

The data that support the findings of this study are openly available in the repository “Dataset for: ‘Disentangling the effects of hearing loss and age on amplitude modulation frequency selectivity’” in DTU Data, at http://doi.org/10.11583/DTU.25134611.

## APPENDIX

### A. Experimental protocol

Table 2 summarizes the experimental protocol used for each HI listener (both young and older). Modulation rate discrimination thresholds were also collected but were not considered in the present study.

**Table 2:** Experimental protocol implemented for all HI listeners, including the tests of hearing threshold at the carrier frequency (HT), modulation rate discrimination (MRD), masked threshold pattern (MTP), and reverse digit span (RDS).

| *Session* | *Test* | *Order* |
| --- | --- | --- |
| 1 | HT followed by<br>MRD | Fixed within<br>session |
| 2-5 | MTP 4 Hz | Random across<br>sessions |
|  | MTP 16 Hz |  |
|  | MTP 64 Hz |  |
|  | MTP 128 Hz<br>followed by AM<br>detection |  |
| Any (2-5) | RDS | Randomly added<br>to any session |

### B. Parameters and post-hoc analyses for masked-threshold patterns

Table 3: summarizes the post-hoc analyses of the ANOVA comparing the MTPs for the older HI and NH listeners (Figure 4).

**Table 3.** Summary of post-hoc analyses of the MTPs comparing older HI and older NH listeners, for each target modulation frequency. The effect sizes (Δ*M*) and standard errors (*SEs*) are reported for each pairwise comparison. Significant p-values are highlighted in bold.

| *Target frequency*<br>*$f_m$ [Hz]* | *Effective masker-modulation bandwidth [Hz]* | *Masker-modulation center frequencies [Hz]* | *Masker-modulation position [octaves re target $f_m$]* | *$\Delta M$ [dB]* | *SE [dB]* | *Corrected p-value* |
| --- | --- | --- | --- | --- | --- | --- |
| 4 | 1.4 | 1 | -2 | -4.6 | 1.1 | **< 0.001** |
|  |  | 1.6 | -4/3 | -3.7 |  | **0.001** |
|  |  | 2.5 | -2/3 | -2.2 |  | **0.047** |
|  |  | 4 | 0 | -1.3 |  | 0.237 |
|  |  | 6.3 | 2/3 | -0.8 |  | 0.472 |
|  |  | 10.1 | 4/3 | -1.6 |  | 0.147 |
|  |  | 16 | 2 | -1.7 |  | 0.134 |
| 16 | 5.6 | 4 | -2 | -4.5 | 0.8 | **< 0.001** |
|  |  | 6.3 | -4/3 | -2.3 |  | **0.003** |
|  |  | 10.1 | -2/3 | -1 |  | 0.196 |
|  |  | 16 | 0 | 0.1 |  | 0.935 |
|  |  | 25.4 | 2/3 | -1 |  | 0.2 |
|  |  | 40.3 | 4/3 | -1 |  | 0.179 |
|  |  | 64 | 2 | -2.9 |  | **< 0.001** |
| 64 | 15.1 | 4 | -4 | -4.6 | 1.2 | **< 0.001** |
|  | 22.3 | 16 | -2 | -3 |  | **0.014** |
|  |  | 25.4 | -4/3 | -2.2 |  | 0.061 |
|  |  | 40.3 | -2/3 | -1.7 |  | 0.152 |
|  |  | 64 | 0 | -1.4 |  | 0.249 |
|  |  | 101.6 | 2/3 | -3 |  | **0.013** |
| 128 | 27.2 | 4 | -5 | -4.9 | 1.2 | **< 0.001** |
|  | 44.6 | 32 | -2 | -3.5 |  | **0.005** |
|  |  | 50.8 | -4/3 | -2.7 |  | **0.027** |
|  |  | 80.6 | -2/3 | -2.2 |  | 0.074 |
|  |  | 128 | 0 | -3.8 |  | **0.003** |

Table 4: summarizes the post-hoc analyses of the ANOVA comparing the MTPs for the older HI and young NH listeners (Figure 5).

**Table 4.**
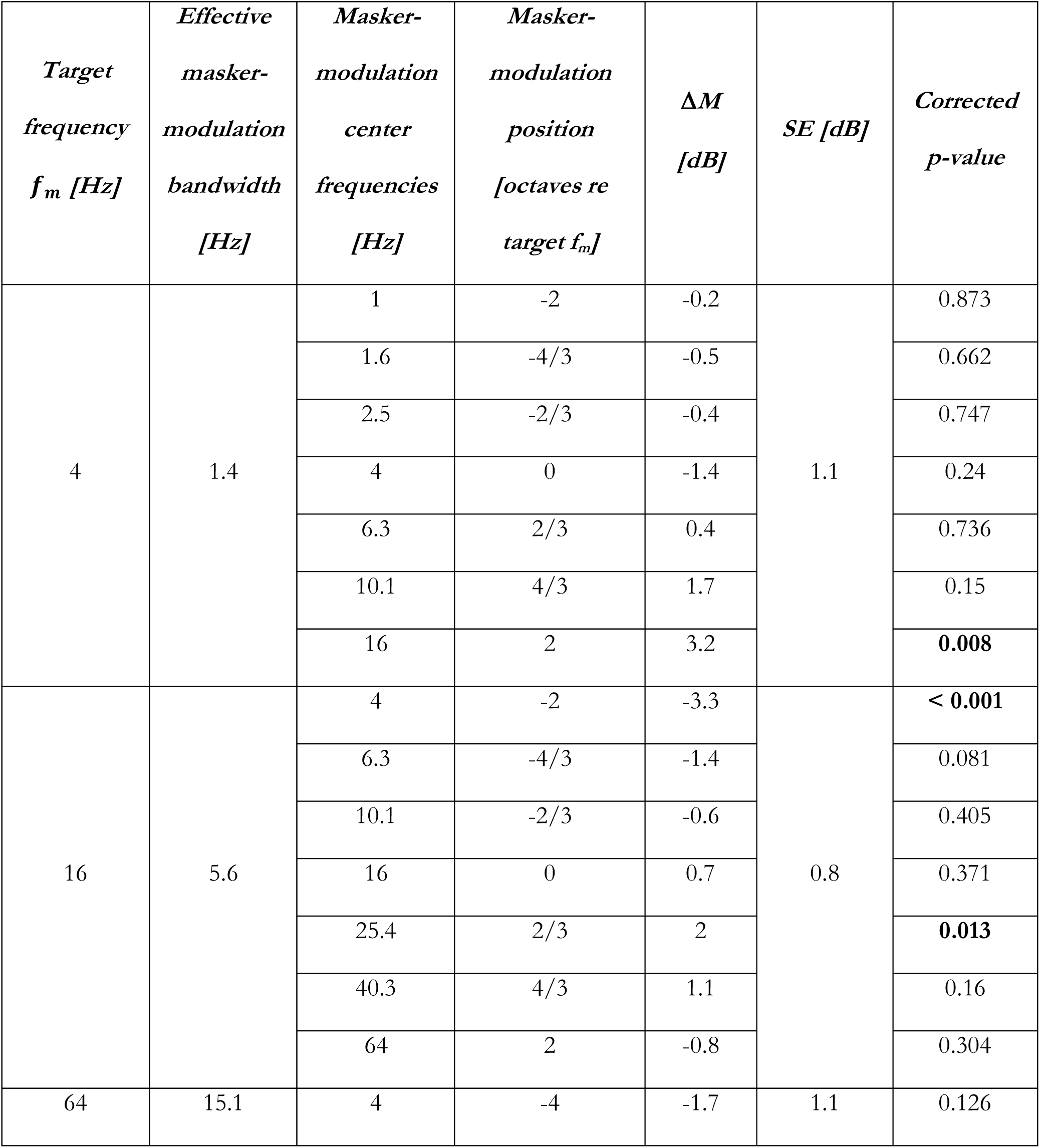
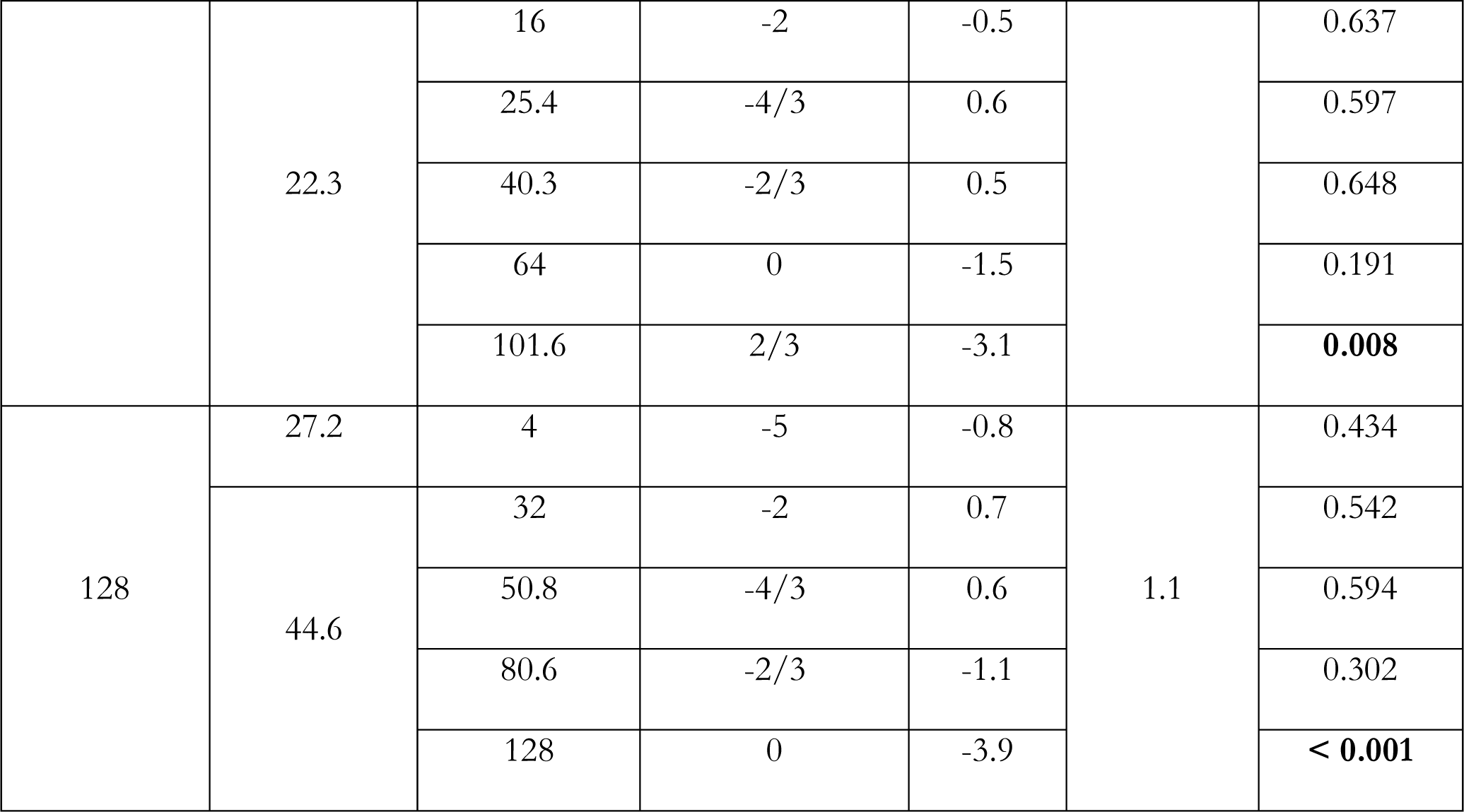
Summary of post-hoc analyses of the MTPs comparing older HI and young NH listeners, for each target modulation frequency. The effect sizes (Δ*M*) and standard errors (*SEs*) are reported for each pairwise comparison. The significant p-values are highlighted in bol

## Footnotes

1 The classification of the collected audiogram as one of the standard audiograms was performed using the lowest mean squared error between the standard audiograms and the audiometric thresholds at frequencies from 1 to 8 kHz.

2 Except for one young HI listener who showed a gap of 20 dB at 4 kHz and one older HI listener who showed a gap of 20 dB at 1 kHz.

3 Overall, 8 measurements were added for 2 of the young HI listeners and 1 measurement was added for a single older HI listener.

4 Overall, a total of 2 measurements were added for a single young HI listener, and 7 measurements for 3 of the older HI listeners, due to aborted measurement runs. Five measurements were added for 2 of the young HI listeners and 10 measurements for 4 of the older HI listeners, due to excessive standard errors. Masked thresholds could not be estimated for a single older HI listener, at masker modulation center frequencies of +2/3 and +4/3 octaves relative to the 4 Hz target modulation frequency.

5 Levene’s test was not passed for one of the models applied, while Shapiro-Wilk’s test was not passed for 3 of the other models. However, the deviations were minor, and the model predictions matched the average group data accurately. Hence, no transformations of the data were applied in order to keep all applied analyses consistent.

