## Supplementary Figure 1 for "Disentangling the effects of hearing loss and age on amplitude modulation frequency selectivity"

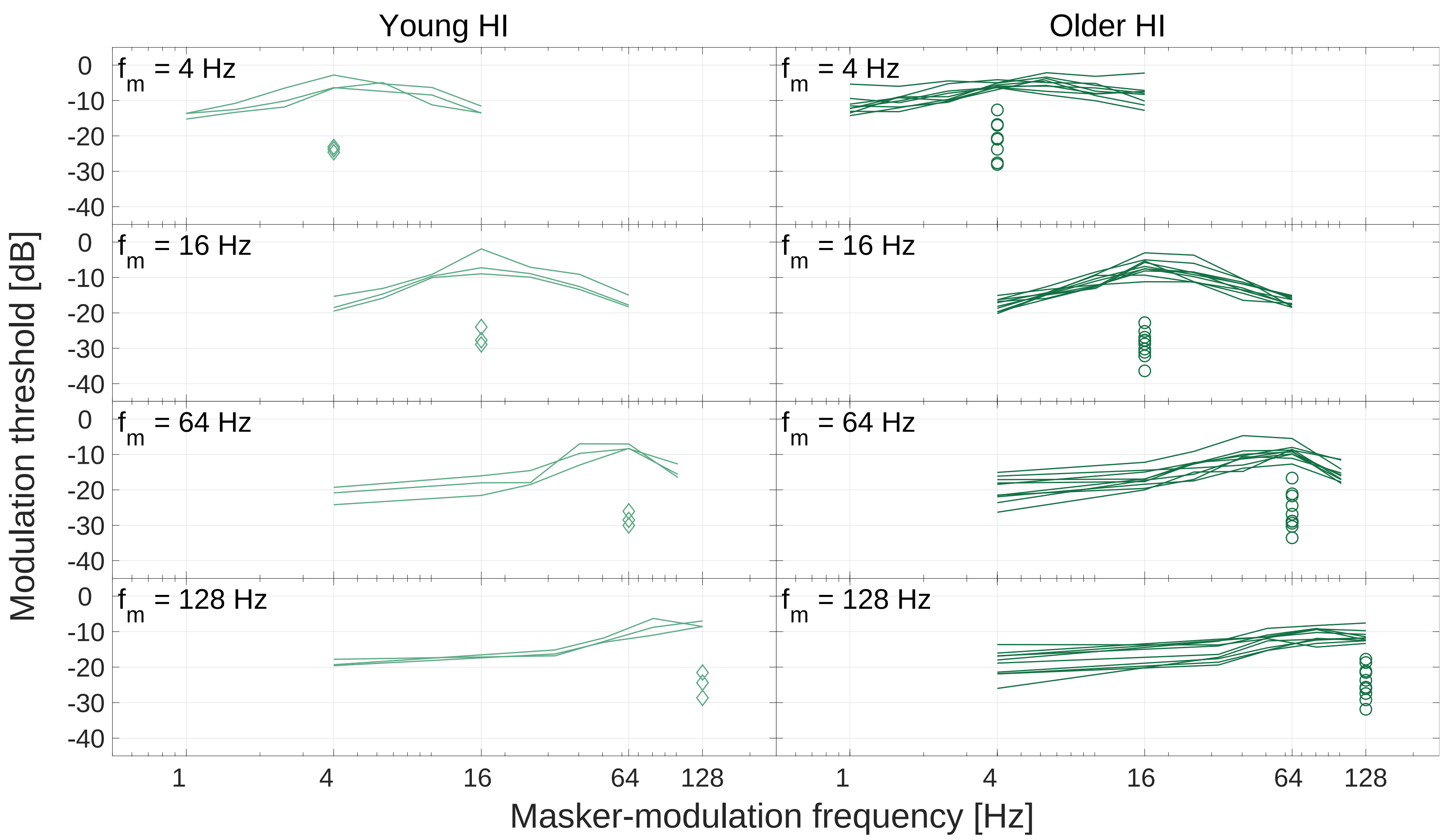


Supplementary Figure 1: Individual MTPs (continuous lines) and AM detection thresholds (open symbols) for all HI listeners, as a function of modulation frequency. The results obtained from young HI (light green lines and diamonds) and older HI listeners (dark green lines and circles) are shown in the left and right columns, respectively. Each row shows the results at a specific target modulation frequency. Note that the data from one of the older HI listeners was excluded from the 4-Hz panel for both the MTP and AM detection threshold.
