## Supplementary Figure 2 for "Disentangling the effects of hearing loss and age on amplitude modulation frequency selectivity"

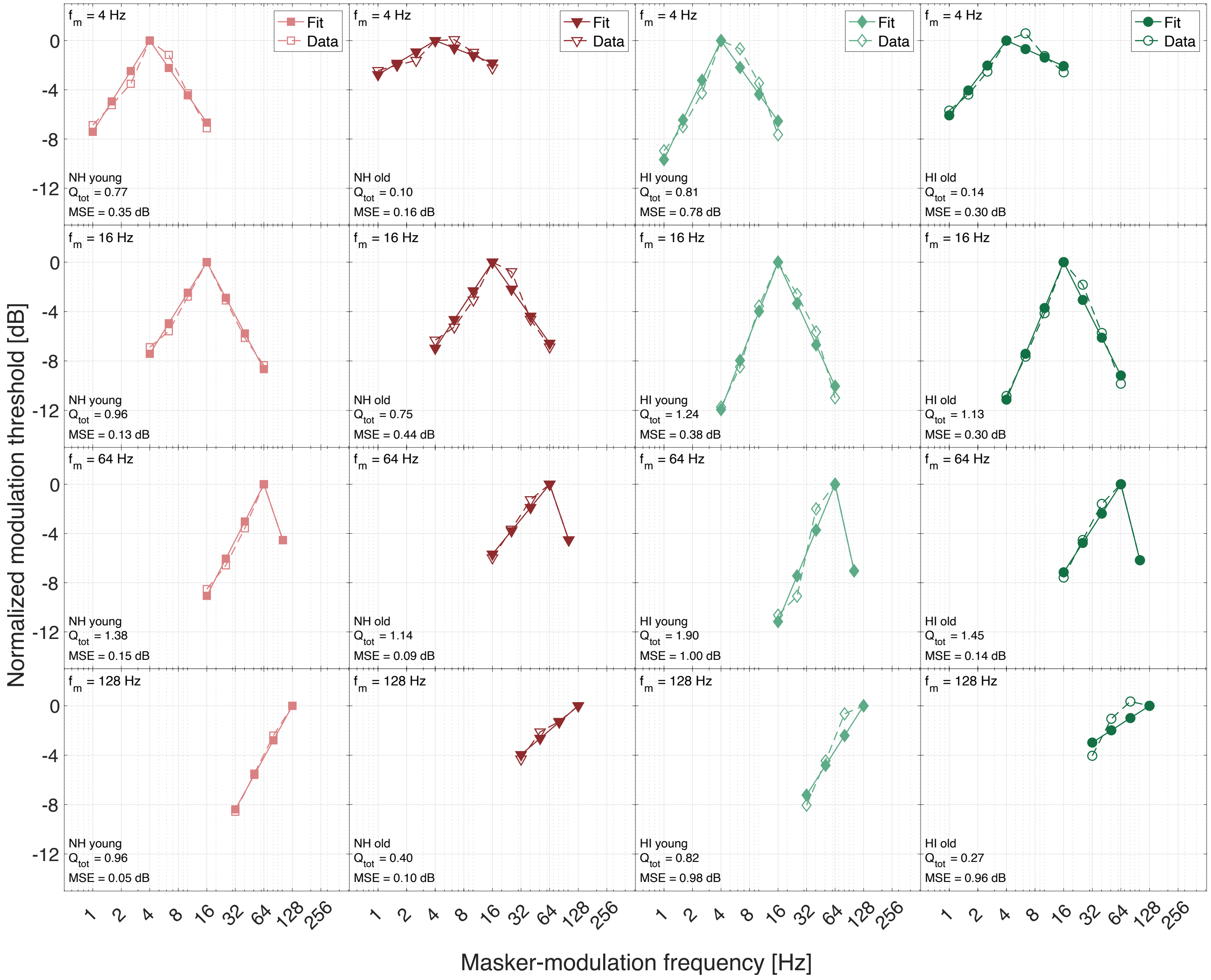


Supplementary Figure 2: Raw thresholds and linear fits for each listener group (columns) and each target modulation frequency (rows). The collected data are shown in open symbols and dotted lines, and the fitted data are shown in closed symbols and continuous lines. The data from young and older NH listeners were obtained by Regev *et al.* (2023) and are publicly available from Regev *et al.* (2024). The thresholds are shown on a normalized scale, i.e., where the on-frequency masked threshold is adjusted to 0 dB.
